# Refate identifies chemical compounds to target trans-regulatory networks for cellular conversion

**DOI:** 10.1101/2025.07.09.664003

**Authors:** Di Xiao, Sonu Sahadevan, Melissa M. Mangala, Hani Jieun Kim, Anna Fredericks, Hao Huang, Raja Jothi, Patrick Tam, Anai Gonzalez-Cordero, Katherine G. Zyner, Pengyi Yang

## Abstract

Identifying chemical compounds that target trans-regulatory networks (TRNs) underlying molecular programs of cells for directed cellular conversion (i.e. differentiation, reprogramming, transdifferentiation, and dedifferentiation) is a key step towards advancing regenerative medicine. Recent innovations in single-cell omics technologies enabled high-resolution profiling of TRNs that govern cell identity and cell-fate decisions. Here, we introduce Refate, a computational framework that integrates large-scale multimodal single-cell atlas data to quantify cell propensity of genes, together with six drug databases, to identify chemical compounds that target TRNs for directed cellular conversion. The reconstructed TRNs, including protein-protein interactions and gene regulatory networks, alongside chemical compounds that drive the cellular conversion provide greater biological interpretability and improve efficiency and efficacy. We evaluated Refate by testing its ability to uncover known transcription factors and chemical compounds validated in experimental conversions of various cell types. Furthermore, we experimentally validated the attribute of several novel chemical compounds identified by Refate for enhancing the conversion of human embryonic stem cells to human cranial neural crest cells. Together, these findings demonstrate Refate as an effective tool for discovering chemical compounds that target TRNs to enable cellular conversion, advancing efforts towards regenerative medicine.

**Highlights:**

- Refate quantifies genes for cellular conversion using multimodal single-cell atlases
- Refate uncovers trans-regulatory networks (TRNs) underlying cellular conversion
- Refate identifies chemical compounds that target TRNs for cellular conversion
- Validation of Refate identified chemical compounds for hESC to hCNCC conversion

## INTRODUCTION

Cell identity and cell-fate decisions are orchestrated by trans-regulatory networks (TRNs) spanning multiple molecular programs, including protein-protein interactions (PPIs), gene regulatory networks (GRNs), and epigenetic regulation. Changing a cell’s identity from one cell type to another, known as cellular conversion, holds great potential for regenerative medicine and has been successfully demonstrated for various cell types through the perturbation of key transcription factors (TFs) and GRNs that underpin cell identity and cell-fate decisions ^1,2^. Depending on the starting and target cell types, cellular conversion can be achieved through experimental approaches such as differentiation, reprogramming, transdifferentiation, dedifferentiation, or combinations thereof. Nevertheless, experimental testing of cellular conversion is highly resource intensive. Developing computational tools to predict molecular drivers for converting cells from a starting to a desired cell type would significantly reduce the time and resources for experimental validation of the functionality of the drivers, and is critical for advancing cellular engineering and regenerative medicine such as endogenous repair ^3^.

Various computational methods have been developed to predict molecular drivers of cellular conversion ^4^. Traditional methods primarily focus on identifying key TFs and GRNs that control cell identity and drive the conversion of cells using predominantly transcriptome profiles ^5^. Notable examples include CellNet ^6^, which predicts TFs for cellular conversion using a compendium of transcriptome profiles compiled from microarray data of diverse tissues and cell types; Mogrify ^7^, which introduces a cell type ontology tree constructed from FANTOM5 expression data; and more recently ANANSE ^8^, which incorporates epigenetic information, such as TF binding profiles and chromatin accessibility from ATAC-seq data, in its predictions; and CellOracle ^9^, which integrates single-cell RNA-sequencing (scRNA-seq) and scATAC-seq data to identify GRNs for cellular conversion. The main outputs from these methods are typically lists of genes and TFs, which still present significant experimental hurdles, requiring labour-intensive TF overexpression to validate and apply predicted drivers for cellular conversion.

To reduce the experimental burden and increase flexibility, a few recent methods aim to predict chemical compounds and small molecules that can modulate TFs and the GRNs they form, thereby drive cellular conversion without TF overexpression or other permanent genetic modifications. Examples include SiPer ^10^, which uses scRNA-seq data of a starting cell type and a compendium of transcriptome signatures to identify candidate compounds for cell conversion, and DECCODE ^11^, which uses expression profiles of cells from the FANTOM5 database and those treated by small molecules in the LINCS project for its predictions. However, SiPer relies on having both scRNA-seq data of the starting cell type and a predefined TF list known to induce the target cell type, limiting its applicability when these resources are unavailable. Meanwhile, DECCODE focuses on matching target cell-type expression profiles, neglects the starting cell type, and only supports conversions among cell types already in its database. Moreover, both methods rely predominantly on transcriptional profiles but overlook epigenetic information such as chromatin accessibility. Developing methods that utilise transcriptional and epigenetic information from large-scale multimodal single-cell omics data, and integrate these with PPIs and GRNs to modulate TRNs, could improve both accuracy and interpretability of chemical compound predictions for directed cellular conversion.

Here, we present Refate, a computational framework designed to identify TRNs and chemical compounds that can drive cellular conversion from a starting cell type to a target cell type. Refate begins by assigning a cell propensity score (CPS) for each gene through integrative analysis of large-scale multimodal single-cell atlas data, which capture both transcriptional and epigenetic landscapes across a large number of human and mouse cell types. For a given pair of starting and target cell types, Refate then identifies potential driver genes and TFs by integrating CPS with the expression differences for individual genes between the start and target cell types. Finally, Refate reconstructs TRNs that underlie the conversion using PPI and GRN databases, and cross-references six drug databases to prioritise chemical compounds that target these TRNs for guiding the intended cellular conversion. Compared to earlier methods, Refate combines transcriptional and epigenetic data at single-cell resolution and enable new cell type conversions to be predicted using only their expression profiles, streamlining the path towards more efficient and versatile cellular engineering.

We benchmarked Refate with other state-of-the-art methods on their ability to unveil chemical compounds that have been experimentally validated for driving the conversion of various cell types. Notably, the multilayered TRNs reconstructed by Refate for cellular conversion offer greater biological interpretability, revealing how predicted chemicals might regulate cell identity. Furthermore, we applied Refate to prioritise chemical compounds that enhance the differentiation of human embryonic stem cells (hESCs) to human cranial neural crest cells (hCNCCs), a valuable system for investigating congenital craniofacial disorders and neurocristopathies ^12^. We subsequently validated several novel compounds predicted by Refate for this conversion. Taken together, these results demonstrate the utility of Refate for characterising TRNs and identifying chemical compounds that target them for directed cellular conversion, bearing significant implications for cell engineering and regenerate medicine.

## RESULTS

### Refate quantifies cell propensity score of genes from multimodal single-cell atlases

The Refate framework, summarised in Figure 1A, first generates a cell propensity score (CPS) for each gene from scRNA-seq and scATAC-seq atlases by computing Cepo statistics, which measure differential stability in gene expression, a key indicator of cell identity ^13^. By integrating Cepo values across cell types from both RNA and ATAC data modalities, the resulting CPS reflects a gene’s capacity to define cell identity. Given a starting and a target cell type, Refate next computes a cellular conversion score (CCS) for each gene by combining CPS with the fold change in gene expression between the two cell types. Refate then constructs TRNs by incorporating GRNs extracted from the TRRUST database (v2) ^14^ and PPIs extracted from the STRING database (2023 version) ^15^. Each TRN is scored based on the individual CCS values of its genes and, finally, these scored TRNs are used as input to prioritise candidate chemical compounds using a combination of six drug databases.

**Figure 1.**
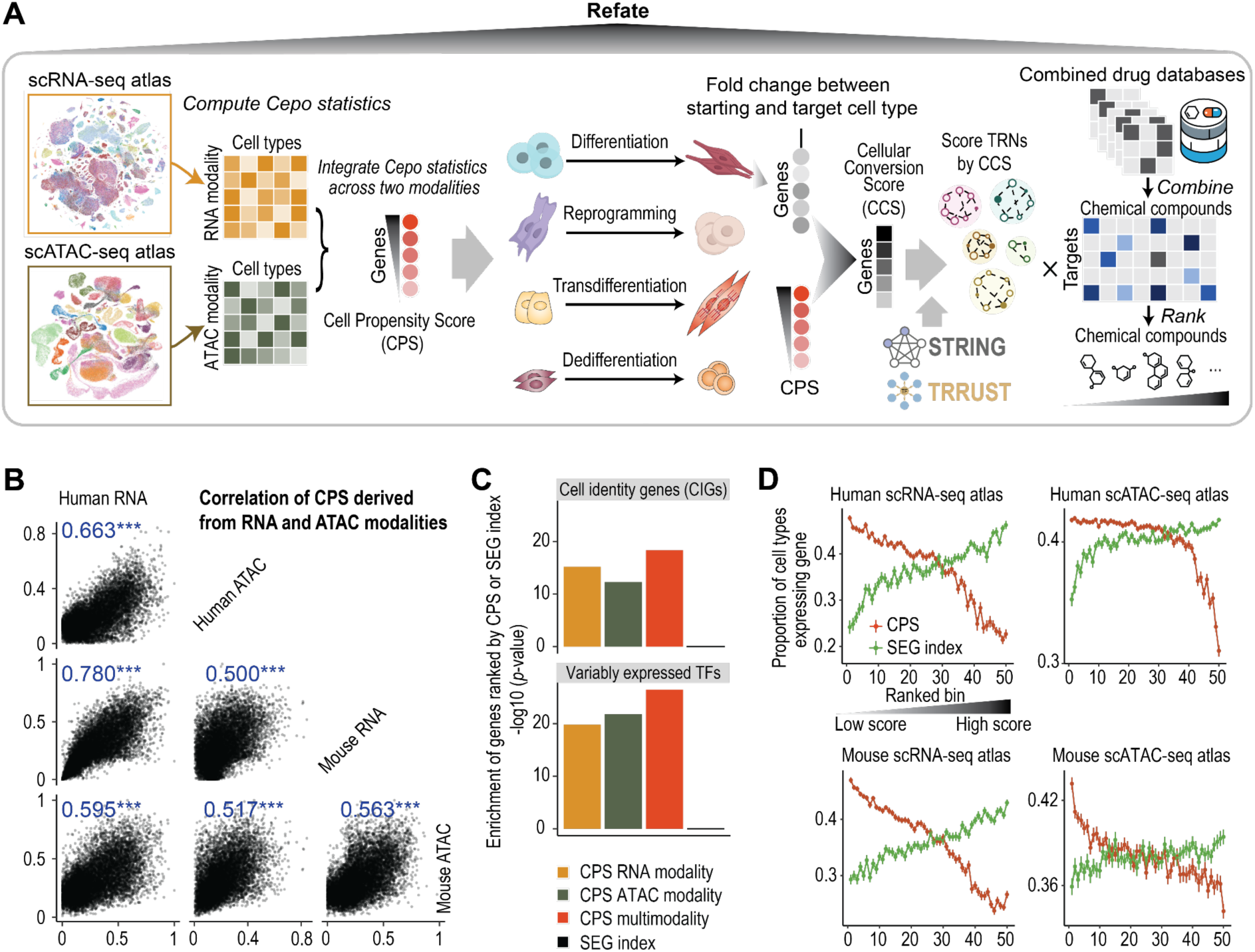
Refate framework for prioritising genes and chemical compounds for cellular conversion. (**A**) Schematic overview of the Refate framework for generating cell propensity score (CPS) of genes, cellular conversion score (CCS) of TRNs, and chemical prioritisation for driving the conversion from a starting and target cell types. (**B**) Correlation analysis of CPS derived from mouse and human scRNA-seq (denoted as mouse RNA and human RNA) and scATAC-seq (denoted as mouse ATAC and human ATAC) atlases. (**C**) Over-representation analysis of gene sets of cell identity genes (CIGs) ^16^ and variably expressed TFs (VETFs) ^17^ using top-ranked genes by CPS derived from data of modality and multimodality. Genes ranked by stably expressed gene (SEG) ^18^ index were included as controls. (**D**) Specificity of gene expression across cell types for genes ranked by CPS derived from multimodal data or by SEG index, respectively. Ranked genes were grouped into 50 bins, and the proportions of the cell types that express the genes in each bin were computed.

To assess if the use of multimodal single-cell data provides complementary information for better identification of genes that affect cell identity, we first calculated the CPS using either RNA or ATAC modality alone using human and mouse scRNA-seq ^19,20^ and scATAC-seq atlases ^21,22^, respectively. We found moderate to strong Pearson’s correlation coefficients, ranging from 0.5 to 0.78, between CPS derived from either RNA or ATAC modalities in the two species (Figure 1B), suggesting that CPS are concordant across the two data modalities and conserved between human and mouse. We next performed enrichment analyses of genes ranked by CPS derived from either RNA or ATAC or their combination using previously and independently defined cell identity genes ^16^ and variably expressed TFs ^17^ (Figure 1C). As a control, we also included the genes ranked by their stable expression gene (SEG) index which marks housekeeping programs of cells ^18^. In both gene sets, we found CPS derived from using combined data modalities resulted in more gene enrichment, suggesting that the integrated analysis of transcriptomic and epigenomic data led to better identification of genes and TFs that underpin cell identity. As expected, genes ranked by the SEG index show no enrichment for the two gene sets.

To further evaluate the CPS of genes for cell identity, we next partitioned the genes into 50 bins based on their scores and then calculated the proportion of cell types that express the genes in each bin. The same analysis was also performed for genes ranked by their SEG index. We found that genes with higher CPS are expressed in a smaller proportion of cell types, indicating their cell type specificity in marking the cell identity programs (Figures 1D, S1A). In comparison, those with higher SEG index tend to be expressed across more cell types for both human and mouse, consistent with their attributes in housekeeping programs. We also assessed the overall expression levels of genes based on their CPS or SEG index (Figure S1B). We found that in general genes with high CPS are expressed at low to moderate levels, consistent with the expression of many cell type specific TFs. In contrast, genes with a high SEG index are expressed at a high level.

To assess the reproducibility of the CPS, we sub-sampled 80% of cells in the human and mouse atlases for calculating CPS repeatedly for 10 times. The high correlation coefficients among CPS calculated from different sub-samples of the full datasets confirm the reproducibility and robustness to data sampling (Figure S1C, D). Together, these analyses suggest that CPS of genes quantified by Refate using multimodal single-cell omics data robustly mark cell identity across human and mouse cells.

### Refate prioritises genes that drive cellular conversion

For a pair of starting and target cell types, Refate computes CCS to prioritise genes for the cell conversion by integrating the gene CPS with the fold change of gene expression between the pair of cell types (see STAR Methods). To test whether Refate can identify driver genes for a cellular conversion of interest, we curated sets of known TFs that were experimentally validated in six different cellular conversions, covering differentiation, reprogramming and trans-differentiation (Supplementary Table 1). We found that CCS calculated from Refate prioritise many of the known TFs across the six cellular conversions (Figure 2A, B), and results from using both data modalities support the identification of these known TFs (Figure S2A). As an example, in differentiating human embryonic stem cells (hESCs) to cardiac muscle cells, known factors such as GATA4, TBX5, and MEF2C ^23^ were among the top genes prioritised by Refate (Figure 2A). Another classical and well described conversion is the reprogramming of fibroblasts to human induced pluripotent stem cells (hiPSCs), where Refate was able to prioritise key reprogramming factors including SOX2, POU5F1 ^24^, MYCN, HMGA1 ^25^, and NR5A2 ^26^ that were experimentally validated for such conversion.

**Figure 2.**
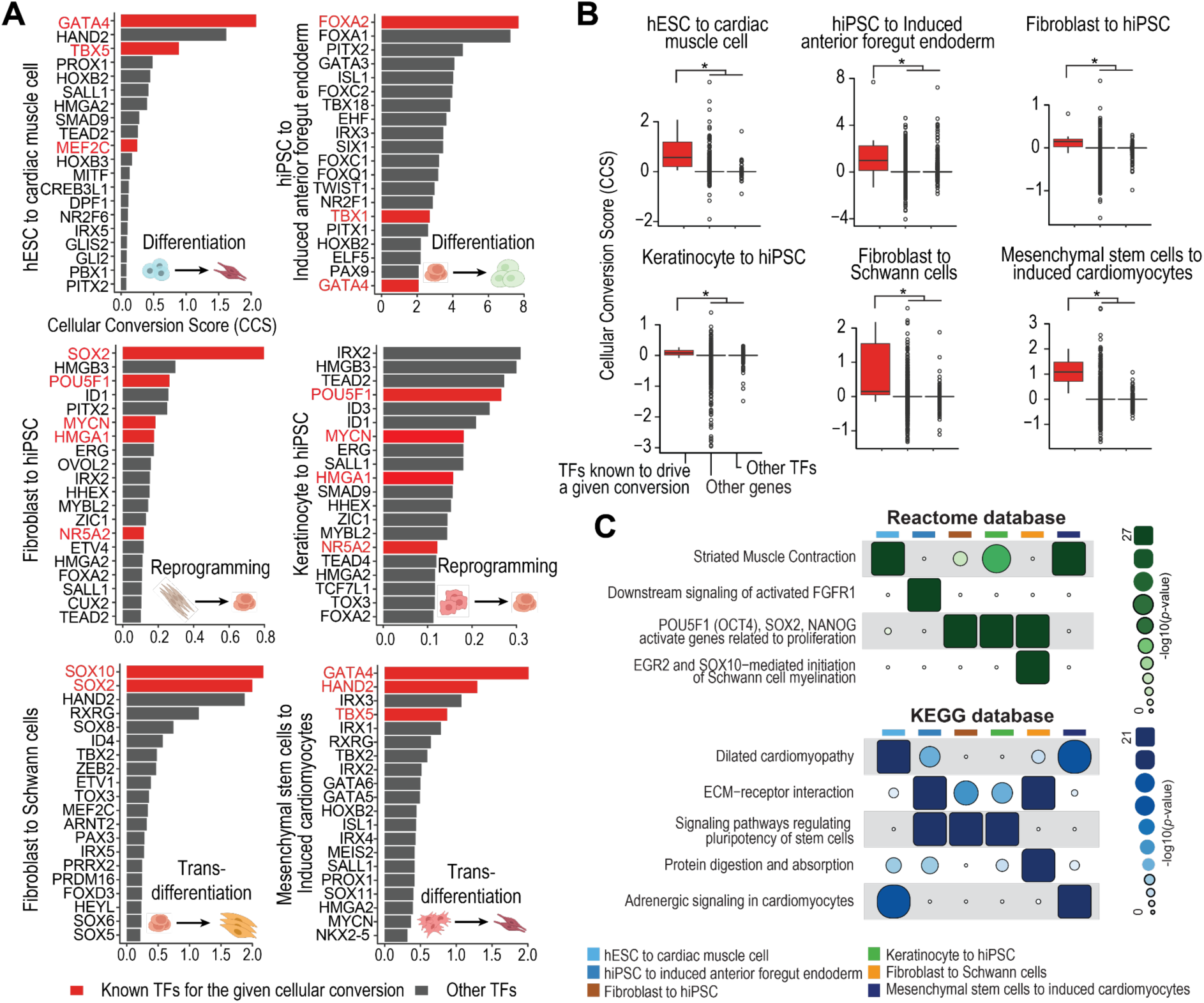
Refate identifies known and novel TFs that drive experimentally validated cellular conversions. (**A**) Barplots showing the top-20 TFs ranked by CCS in each of the six experimentally validated cellular conversions of differentiation, reprogramming, and trans-differentiation. Known TFs for their respective cellular conversions are highlighted in red. (**B**) Boxplots of CCS for known TFs for cellular conversion, other TFs, and all other genes. * denote statistical significance from a two-sided Wilcox rank sum test (p<0.05). (**C**) Most enriched pathways of the CCS top-ranked genes for the six different cellular conversion using Reactome and KEGG databases.

Beyond known TFs, Refate identified numerous novel candidate genes for their respective cellular conversions. We computationally assessed these gene sets by performing pathway over-representation analyses using the top-ranked genes from each cellular conversion against the Reactome and the KEGG databases (Figure 2C, S2B). For example, the pathway related to the reprogramming factors of POU5F1, SOX2, and NANOG were highly enriched in hiPSC reprogramming. The pathway striated muscle contraction involved in cardiac muscle cell formation ^27^ was highly enriched in hESC to cardiac muscle cell differentiation (Figure 2C). These results affirm the cell type specificity of gene sets prioritised by Refate with respect to their target cell types. Taken together, these findings demonstrate that Refate identify both previously known and potential novel drivers for targeting cellular conversion.

### Refate leverages multiple drug databases to prioritise chemical compounds for cell conversion

To prioritise chemical compounds that target a specific cellular conversion, Refate first constructs TRNs that span protein complexes and gene networks by integrating PPIs and GRNs extracted from STRING and TRRUST databases. Second, it partitions them into sub-networks for quantification of their impact on the cellular conversion using the Louvain community detection algorithm and CCS of genes contained in the sub-networks (STAR Methods). Next, Refate scores a chemical compound for a desired cellular conversion by quantifying and summarising its effect on each of all the sub-networks based on the genes in the sub-networks that are targeted by the chemical compound. To achieve high coverage on chemical compounds and their targets, six drug databases including CTD ^28^, DGIdb ^29^, DrugBank ^30^, DrugRepurpose ^31^, STITCH ^32^ and TTD ^33^ are curated in Refate (Figure S3A), and the direction and evidence of each chemical compound-target relationships are considered and weighted (STAR Methods).

By evaluating a large collection of chemical compounds that have been experimentally validated for 18 cellular conversions covering various cell types (Supplementary Table 2), we found that, overall, Refate performed significantly better on prioritising chemical compounds that are known to drive cell conversions compared to SiPer and DECODE, two state-of-the-art methods for identifying chemical compounds for cellular conversion (Figure 3A, S3B). A better performance and high coverage of Refate are further demonstrated in each cellular conversion where, in most cases, Refate better prioritised experimentally validated chemical compounds in converting cell types compared to the two alternative methods (Figure 3B). Since SiPer also relies on a curated drug database for chemical compound identification for cellular conversion, we explored the utility of the database curated by SiPer in our Refate framework (Figure S3B). Our analysis revealed that using the six chemical compound-target databases curated for Refate resulted in improved prioritisation of experimentally validated chemical compounds, both overall (Figure S3C) and across cell types (Figure S3D).

**Figure 3.**
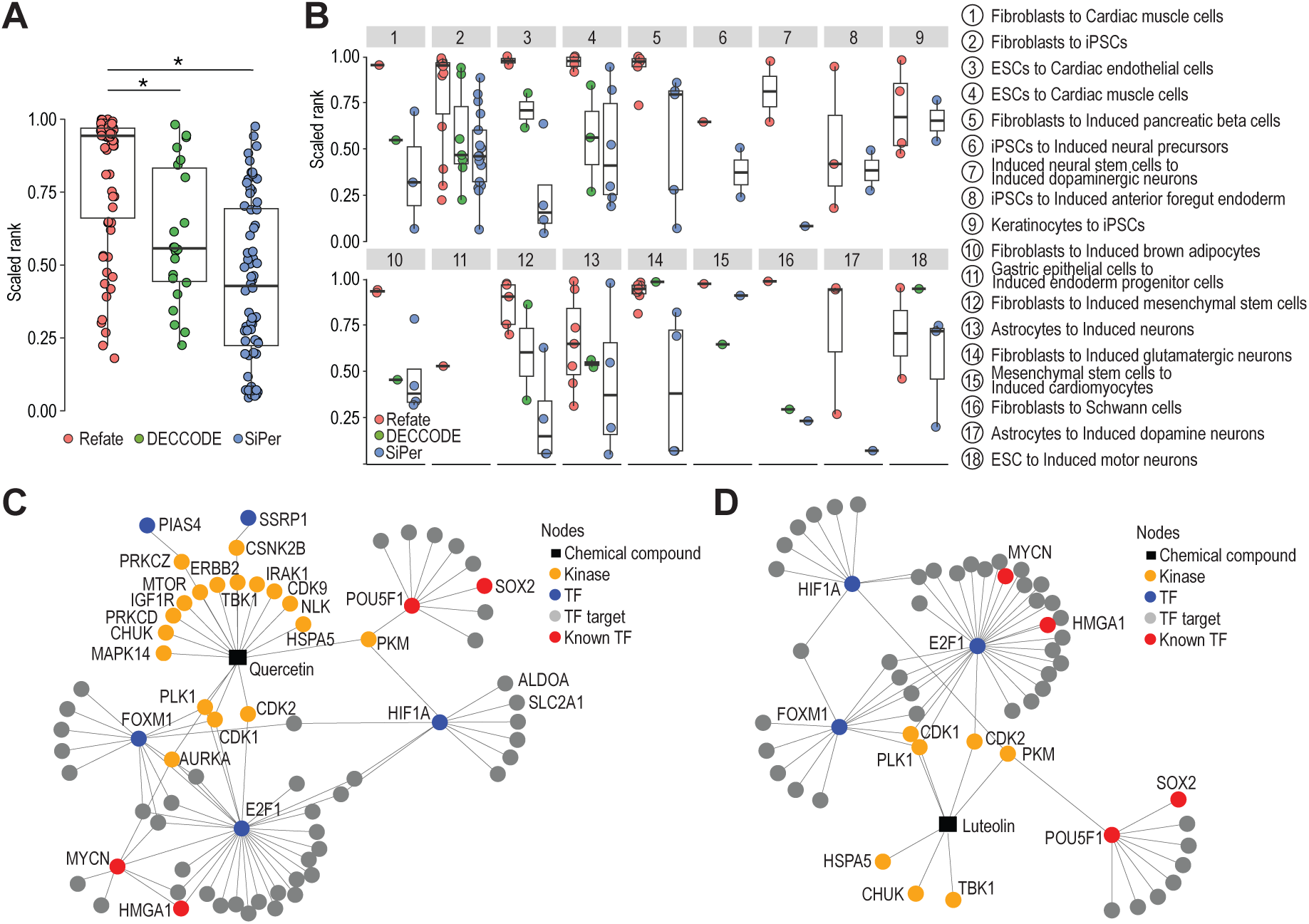
Refate prioritises known and novel chemical compounds for various cellular conversions. (**A**) Boxplots showing the overall scaled rank of experimentally validated chemical compounds for 18 cellular conversions by either Refate, DECCODE, or SiPer. * denote statistical significance from a two-sided Wilcox rank sum test (p<0.05). (**B**) Boxplots showing the scaled rank of known chemical compounds for each of the 18 curated cellular conversions by either Refate, DECCODE, or SiPer. (**C**) Reconstruction of TRN targeted by Quercertin, a known compound, for fibroblast to iPSC reprogramming. (**D**) Reconstruction of TRN targeted by Luteolin, another compound prioritised by Refate for fibroblast to iPSC reprogramming. Nodes represent genes and edges denote interactions curated in drug, PPI, and TF-target databases.

To elucidate the effects of chemical compounds on cellular conversion, we reconstructed the TRNs to unveil how these compounds interact with key regulators to mediate a given conversion. For example, the TRN reconstructed for Quercetin, a top-ranked chemical known to drive and enhance fibroblast to iPSC reprogramming ^34^, revealed its interactions with (i) PKM and HIF1A, suggesting regulation of glycolytic metabolism and hypoxic adaptation; (ii) CDK1/2 and E2F1, indicating roles in cell cycle progression; (iii) MYCN and HMGA1, associated with proliferation and chromatin remodelling; and notably, (iv) POU5F1 and SOX2, implicating direct regulation of pluripotency networks (Figure 3C). Similarly, the TRN reconstructed for Luteolin, another predicted compound for enhancing fibroblast to iPSC reprogramming, exhibited many of the same interactions identified for Quercetin, reinforcing its potential role in pluripotency regulation (Figure 3D). Collectively, these results highlight Refate’s utility for identifying chemical compounds to target TRNs to drive cellular conversion.

### Experimental validation of Refate predicted chemical compounds for hESC to hCNCC differentiation

hCNCCs contribute to the formation of a diverse range of craniofacial tissues, making them a valuable *in vitro* model for studying congenital craniofacial disorders and neurocristopathies ^12^. Additionally, hCNCCs provide a source for generating patient specific craniofacial structures and cells, facilitating applications in personalised and regenerative medicine ^35^. While previous studies have established the protocol for differentiating hESCs to SOX10-expressing hCNCCs, the differentiation efficiency remains suboptimal, typically ranging from 40 to 60% ^36,37^. To enhance differentiation efficiency, we applied Refate to this cellular conversion and experimentally evaluated the effects of six compounds selected from Refate’s top prediction list, including LDN193189, XL147, SB203580, Melatonin, Rimonabant, and Kenpaullone (Figure 4A, STAR Methods).

**Figure 4.**
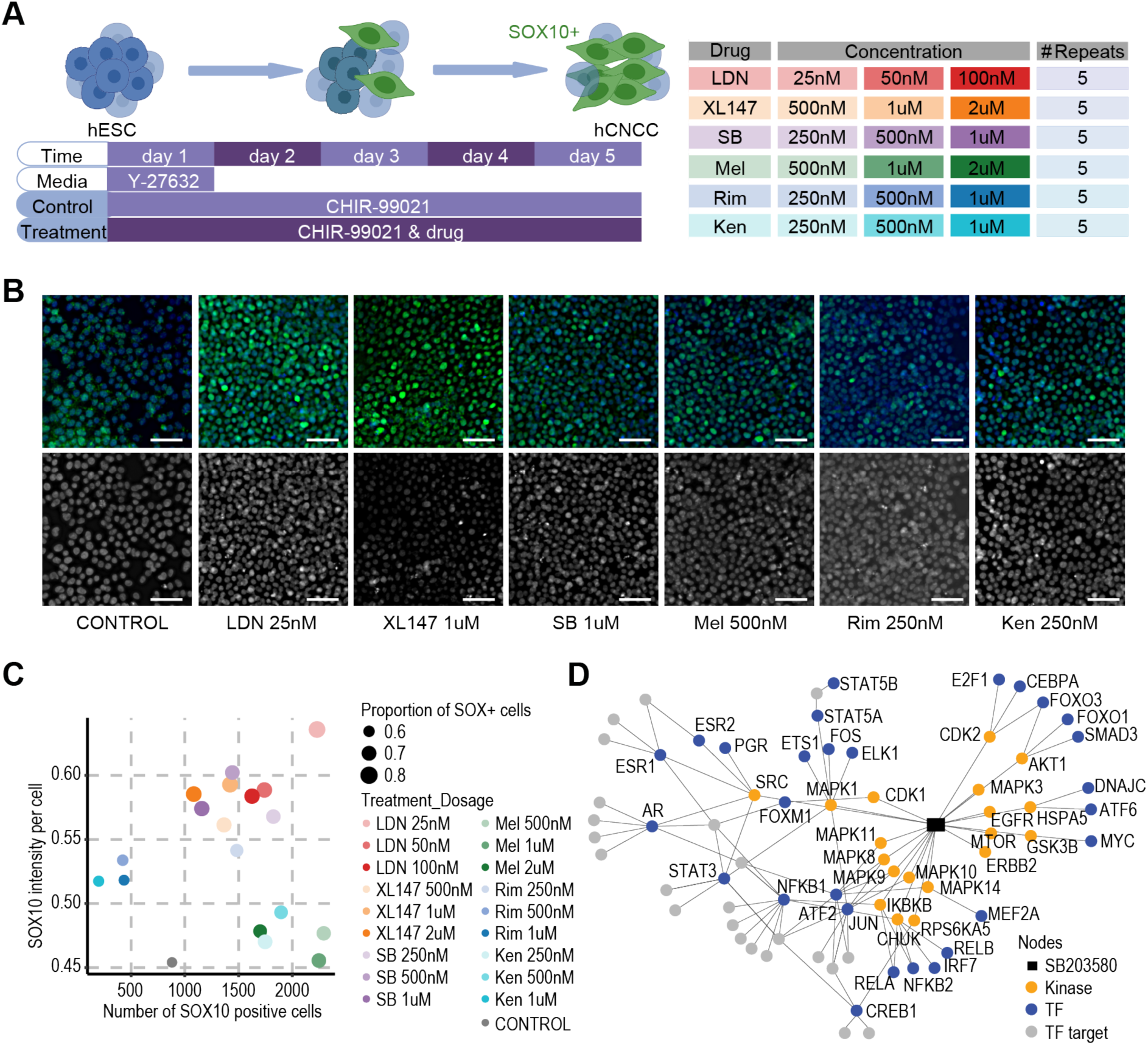
Experimental validation of Refate-predicted small molecules for enhancing hESC to hCNCC differentiation. (**A**) Schematic representation of the experimental design. hESCs were differentiated into hCNCCs using the standard five-day CHIR-99021-based differentiation protocol, with or without the addition of predicted small molecules. Six small molecules were tested at three different concentrations to evaluate their potential to enhance hCNCC differentiation. (**B**) Representative fluorescence microscopy images of hCNCCs stained with SOX10 (green) and DAPI (blue) (upper panel), along with corresponding DAPI images of the cells (lower panel). Scale bars represent 50 µm. (**C**) Scatter plot summarising the effects of chemical compound treatments on hCNCC differentiation based SOX10 marker expression. The x-axis represents the number of SOX10-positive cells, and the y-axis represents the SOX10 intensity per cell. The size of each dot corresponds to the proportion of SOX10-positive cells. (**D**) TRN reconstructed for SB (SB203580), a novel compound prioritised by Refate for driving hESC to hCNCC differentiation. Nodes represent genes and edges denote interactions curated in drug, PPI, and TF-target databases.

Consistent with the standard protocol, the control condition achieved an hCNCC differentiation efficiency around 60%, and several compounds significantly enhanced differentiation efficiency compared to the control (Figure 4B,C and Figure S4A-D). Notably, different compounds exhibited distinct effects on the differentiation of hESCs to hCNCCs relative to the control. Treatment with LDN193189, XL147, and SB203580 led to an increase in both the proportion of SOX10-positive cells and the SOX10 intensity per cell, indicating an overall enhancement in differentiation. In contrast, Rimonabant increased SOX10 intensity per cell without substantially raising the proportion of SOX10-positive cells, suggesting it enhances SOX10 expression levels in individual cells rather than expanding the overall population of SOX10-expressing cells. Melatonin, on the other hand, increased the proportion of SOX10-positive cells while maintaining SOX10 intensity per cell at levels comparable to the control, suggesting that it promotes a broader induction of SOX10 expression across the cell population rather than enhancing per-cell expression levels.

TRN reconstruction offers further insight into potential drug mechanisms. For example, SB203580 targets multiple p38 MAPK isoforms (e.g., MAPK8/9/10/11/14) involved in neural crest migration and differentiation ^38^. Its interactions with TFs such as STAT3, JUN, NFKB1, and RELA suggest a role in early lineage commitment ^39–42^, while connections to FOXO1/3 and CEBPA indicate possible involvement in metabolic regulation and oxidative stress response during differentiation ^43,44^. Additional connections to EGFR, AKT1, and MTOR highlight potential impacts on survival and differentiation ^45,46^, with possible crosstalk to Wnt/BMP pathways via MAPK-associated factors (FOS, ELK1, STAT5A) (Figure 4D). Altogether, these results provide potential avenues for optimising hCNCC differentiation protocols and highlight the utility of Refate for guiding experimental validation for cellular conversions.

## DISCUSSION

Substantial effort has been made to curate molecular profiles and databases for identifying and quantifying genes that drive cellular conversion ^6,7^. Nevertheless, the reliance on ‘bulk’ expression data that profile large cell populations remains a key challenge in deciphering genes that underpin cell identity and cell-fate decisions. The recent development of single-cell omics technologies provides a new opportunity to quantify gene variability within and across a large number of cell types that could not be obtained from bulk data. Leveraging the large-scale multimodal single-cell omics data of a wide range of human and mouse cell types, Refate computes cell propensity scores of genes and subsequently uses this information to quantify genes for their potential in driving any cellular conversion. The increasing availability of single-cell omics data with additional data modalities and from more cell types, tissues, and species will further improve the identification and quantification of genes that may influence the desired cellular conversion.

For a given cellular conversion, many known TFs are prioritised by Refate on the basis of high CCS. Yet, there are many more TFs and other non-TF genes that also exhibit high CCS (Figure 2A,B). While some of these may act as “passengers” rather than drivers of cellular conversion, others could represent unidentified co-factors that complement and contribute to the GRNs and PPIs that drive the process. A key advantage of Refate’s CCS calculation and TRN approach is its ability to integrate the individual effects of TFs and other genes. Indeed, prioritising combinations of genes can lead to more efficient and efficacious cell conversion than enforcing the expression of single TFs which may result in the incomplete cellular conversion of cells ^47^. The TRN-guided prioritisation of chemical compounds by Refate consolidates combinatorial effects of genes and can significantly reduce the complexity of experimentally validating large combinations of TFs and genes in GRNs.

It is worth noting that several recent high-throughput perturbation studies (e.g. over-expression) have been conducted to screen for transcription factors that induce cell-fate conversion and directed differentiation ^48–50^. Due to the scale of these experiments, most studies have focused primarily on evaluating TFs for their role in cellular conversion, while largely excluding non-TF genes and chemical compounds as potential modulators. However, further development of computational analytics could integrate these perturbation data into Refate for in-depth TF identification and TRN reconstruction and more precise prioritisation of compound prioritisation for cellular conversion. Further advancements in computational analytics could enable the integration of perturbation data into Refate, facilitating more comprehensive TF identification and TRN reconstruction, and improved prioritisation of chemical compounds for enhancing cellular conversion.

### Limitations of this study

While most drug databases provide information on the direction of effects for chemical compounds on their target genes, a significant proportion still lack this detail. As a result, Refate may prioritise compounds as candidates even if they act as negative regulators in a given cellular conversion. This limitation is expected to improve as more comprehensive data on chemical effects become available in these databases. Another shortcoming of Refate is its inability to quantitatively predict the optimal dosage and timing of chemical compound administration during cellular conversion. Lastly, the method may prioritise a large number of chemical compounds and the selection of the most suitable ones for functional evaluation remains a challenge. Therefore, effective validation still requires substantial prior knowledge, such as the general effects of a compound and a specific cellular conversion. Therefore, substantial prior knowledge, such as the general effects of compounds and a specific cellular conversion, are still required for selecting the prioritised chemical compounds for validation.

## ACKNOWLEDGEMENTS

This work is supported by a National Health and Medical Research Council Investigator grant (1173469) and a Metcalf Prize to P.Y.; A Children’s Medical Research Institute Postdoctoral Fellowship Award to K.G.Z.; Intramural Research Program of the National Institutes of Health, National Institute of Environmental Health Sciences (1ZIAES102625) to R.J; and the Rebecca Cooper Foundation, through an Al & Val Rosenstrauss Fellowship to A.G.-C.

## AUTHOR CONTRIBUTIONS

Conceptualisation, D.X., K.G.Z. and P.Y.; investigation and formal analysis, D.X., S.S.M.K., K.G.Z. and P.Y.; method development: D.X., H.J.K., H.H., A.F., R.J. and P.Y.; data generation, S.S.M.K., M.M., A.G.-C. and K.G.Z.; writing - original draft, D.X., S.S.M.K., K.G.Z and P.Y.; writing - review & editing, all authors; supervision, P.P.L.T., A.G.-C., K.G.Z. and P.Y.; funding acquisition, A.G.-C., K.G.Z. and P.Y.

## STAR METHODS

KEY RESOURCES TABLE

| REAGENT or RESOURCE | SOURCE | IDENTIFIER |
| --- | --- | --- |
| **Software and algorithms** |  |  |
| Refate 1.0 | This work | https://github.com/PYangLab/Refate |
| R version 4.3.3 |  | https://www.r-project.org/ |
| PhosR 1.13.1 | Kim et al. <sup>51</sup> | https://bioconductor.org/packages/release/bioc/html/PhosR.html |
| Cepo 1.11.2 | Kim et al. <sup>13</sup> | https://github.com/PYangLab/Cepo |
| NetworkD3 | Allaire et al. <sup>52</sup> | https://CRAN.R-project.org/package=networkD3 |
| SiPer | Zheng et al. <sup>10</sup> | https://git-r3lab.uni.lu/menglin.zheng/SiPer |
| DECCODE | Napolitano et al. <sup>11</sup> | https://fantom.gsc.riken.jp/5/cellconv/ |
| Fiji | Schindelin et al. <sup>53</sup> | https://imagej.net/software/fiji/ |
| CellProfiler | Stirling et al. <sup>54</sup> | https://cellprofiler.org/ |
| **Small molecule treatments** |  |  |
| CHIR99021 | Tocris | Cat. No. 4423 |
| LDN193189 hydrochloride | Sigma-Aldrich | SML0559 |
| XL147 | Cayman Chemical | 18011 |
| SB203580 | Cayman Chemical | 152121-47-6 |
| Melatonin | Sigma-Aldrich | M5250 |
| Rimonabant | Cayman Chemical | 168273-06-1 |
| Kenpaullone | SelleckChem | S7917 |
| Y-27632 | StemCell<br>Technologies | 72304 |
| **Experimental models: Human cell lines** |  |  |
| H9 (human embryonic stem cells, hESCs) | WiCell | WA09 |
| **Culture media and reagents** |  |  |
| mTeSR Plus | StemCell<br>Technologies | 100-0276 |
| Matrigel-coated plates | Corning | 354277 |
| Versene | Life Technologies | 15040066 |
| StemPro Accutase Cell<br>Dissociation Reagent | Life Technologies | A1110501 |
| DMEM/F12 | Life Technologies | 10565018 |
| Serum-free B27 supplement | Life Technologies | 17504044 |
| Penicillin-Streptomycin | Life Technologies | 15140122 |
| **Immunohistochemistry reagents** |  |  |
| Paraformaldehyde | Scharlau | 30525-89-4 |
| Triton X-100 | Sigma | T8787 |
| Bovine Serum Albumin (BSA) | Merck | A3983 |
| Tween-20 | Amresco | 0777 |
| DAPI | ThermoFisher Scientific | 62247 |
| Ibidi Mounting Media | Ibidi | 50001-4 |

## RESOURCE AVAILABILITY

### Lead contact

### Materials availability

This study did not generate new unique reagents.

### Data and code availability

Human scRNA-seq data from the Tabula Sapiens Consortium ^19^, human adult scATAC-seq data from the human scATAC-seq atlas ^21^, mouse scRNA-seq data from The Tabula Muris Consortium ^20^ and the mouse scATAC-seq atlas ^22^. The Refate R package and associated vignette are available from Github repository at https://github.com/PYangLab/Refate. Shiny app is available at http://shiny.maths.usyd.edu.au/Refate_Shiny/.

## METHOD DETAILS

### Processing of human and mouse multimodal single-cell omics data

For the human scRNA-seq atlas data, we utilised the pre-processed data generated from the Tabula Sapiens project. In particular, the data were pre-processed to exclude genes encoded by the mitochondrial genome and eliminate cells with less than 200 detected genes or less than 2500 unique molecular identifiers (UMIs). Genes not expressed in any cells were filtered in the data. The pre-processed human scRNA-seq atlas contains 38,203 genes from 437,249 cells annotated to 177 distinct cell types ^19^. Similarly, the pre-processed data for the mouse scRNA-seq atlas was downloaded from the Tabula Muris project. In this dataset, cells were excluded if they had fewer than 500 detected genes or fewer than 1,000 UMIs. This filtering led to the pre-processed mouse scRNA-seq atlas containing 19,817 genes from 55,656 cells annotated to 56 different cell types ^22^.

Gene activity scores were obtained for both human and mouse scATAC-seq atlas data from their respective original studies and a logarithmic transformation was conducted on these scores. In particular, the human scATAC-seq atlas used chromatin accessibility at the promoter as a proxy for gene activity. The promoter was defined as Reads Per Million (RPM) of +/-1 kilobase (kb) around the transcription start site (TSS). Following this process, there are 60,286 genes from 615,998 cells, annotated to 111 cell types ^21^. The mouse scATAC-seq atlas aggregated information across all differentially accessible sites linked to a target gene to compute the “gene activity score” using Cicero ^55^. This procedure led to 20,783 genes from 81,173 cells, annotated to 222 cell types ^22^.

### Processing of additional transcriptomics data

To validate the Refate framework on prioritising TFs for given cellular conversions, we also included additional datasets that captured the transcriptional profiles of human ESCs and iPSCs using scRNA-seq ^56,57^, respectively. In particular, the scRNA-seq data were averaged across cells to obtain pseudo-bulk expression profiles of the cell types. These data in the count scale were first log-transformed and then quantile normalised with respect to human scRNA-seq atlases.

### Cell propensity score (CPS) of genes

We propose CPS as a quantification of how likely a gene is to affect the identity of cells. We derive CPS by first calculating the Cepo statistics ^13^, which uses differential stability to measure genes for marking cell identity, across human and mouse scRNA-seq and scATAC-seq atlases, respectively. Let us denote Cepo statistic of a gene *g* for a cell type *i* as *ds^g^_i_* and *ds* = 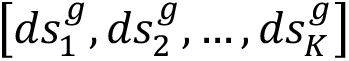 denotes the Cepo statistics of *g* in all *K* cell types in an atlas, the CPS for *g* is defined as 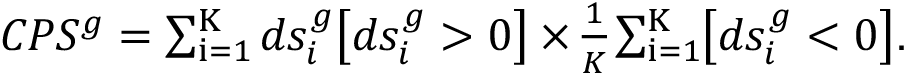. The first component 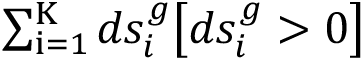 sums the positive Cepo statistics across the cell types with a higher value indicating *g* being a cell identity gene across one or more cell types. The second component 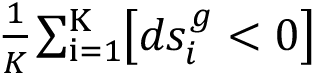 quantifies the proportion of cell types in which *g* does not mark their identity. The two components together prioritise highly cell type specific identity genes that are likely to affect the conversion of cell types. Let us further denote CPS of *g* derived from RNA and ATA 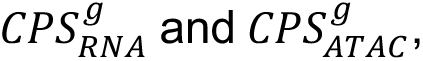 respectively, and the combined CPS of the two data modalities can be defined as 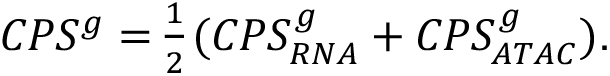

### Cellular conversion score (CCS) of genes

For a given conversion from cell types *a* to *b*, Refate computes CCS for a gene *g*, denoted as 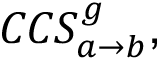 by first calculating the log2 fold change of the expression of *g* between the starting (*a^g^*) and target (*b^g^*) cell types and then multiplying this by CPS of the gene 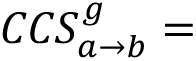 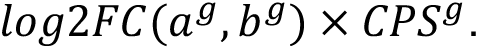 By combining the fold change in expression of a gene in the starting and target cell types and its general potential in governing cell identity as quantified by CPS, Refate prioritises genes that are promising candidates for driving the cellular conversion of interest.

### Construct TRNs and prioritise chemical compounds

For a given cellular conversion, Refate constructs TRNs and ranks chemical compounds based on their potential effects on these TRNs. Specifically, to build TRNs, we first construct an undirected global graph using the igraph package ^58^, integrating PPI data from the STRING database and the GRN data from the TRRUST database. Next, utilising the Louvain community detection method, we identify community structures within TRNs and partition the primary TRNs into distinct and inter-connected sub-networks. To prioritise chemical compounds, we first curated an integrated chemical compound-target database by combining six different drug databases, including DGIdb 4.0 ^29^, DrugBank ^30^, Drug Repurposing hub ^31^, STITCH ^32^, TTD ^33^ and CTD ^28^. To quantify the evidence of each chemical compound-target pair across databases, we took the union of chemical compound-target pairs from all drug databases and then weighted each pair by *log*2(*n* + 1) where *n* is the number of drug databases reporting a given chemical compound-target pair. To account for the directionality of the chemical compound-target interactions, we integrated directional information sourced from the six drug databases. For the chemical compounds that have a negative impact on the targets (description includes keywords such as “Inhibition”, “Decrease”, “Suppressor”, “Inactivator”, “Disrupter”, “Degrader”, “Breaker”, “Antagonist”), we applied a negative sign to the weights derived in the preceding step. Then, we prioritised chemical compounds based on their effects on TRNs. This was achieved by quantifying their influence on the sub-networks identified by the Louvain algorithm and then integrating this evidence into an overall effect. In particular, for a given cellular conversion, after community detection, we derive *m* sub-networks each denoted as *S(i*) where *i* = 1,2,…, *m*. When a drug targets a specific sub-network *S(i*), we compute an individual effect score for each gene in that sub-network as 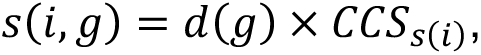 where *g* represents gene that is targeted by a given chemical compound in sub-network *S*(*i*), *d(g)* represents the drug effect on gene *g* and *CCS*_s(i)_ represents the summed CCS for sub-network *S*(*i*). We then aggregate the drug effect *e(i)* on a given sub-network *S*(*i*) by summing all positive scores 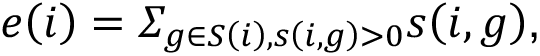 and finally, compute the overall effect of a drug on a cellular conversion by summing the effects over all sub-networks 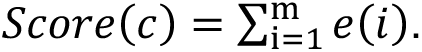 For comparison with other methods, the overall scores from all chemical compounds were ranked and scaled for each tested cellular conversion.

### Chemical compound prediction using SiPer and DECCODE

DECCODE and SiPer pipelines were included for comparison for prioritising chemical compounds. In particular, SiPer pipeline predicts cellular conversions based on a scRNA-seq dataset profiled from the starting cell type and a curated set of transcription factors (TFs) that are responsible for converting cells from the starting cell type to the target cell type ^10^. To run SiPer pipeline, we first prepared scRNA-seq data of the starting cell types and sets of TFs that are reported in the literature for driving a given cellular conversion. The SiPer pipeline was applied to generate Jaccard indices, which prioritise chemical compounds based on the similarity between their target proteins and predicted signalling proteins. To enable comparison of the scores from different methods, the resulting Jaccard indices of all chemical compounds from SiPer were ranked and scaled as was performed for Refate.

For DECCODE, it first transforms the pre-and post-perturbation gene expression profiles (DEPs) in the CMap database ^59^ into differential pathway-based expression profiles (DPEPs). Subsequently, a search is conducted within the CMap database to identify chemical compounds that exhibit DPEPs similar to the query profile. To run DECCODE, we first calculated the log2 fold change of the target cell types against the background of all other cell types in the human scRNA-seq atlas according to the original study ^11^, and converted these gene-based profiles to pathway-based profiles using the gep2gep package ^60^. Next, we calculated the similarity between the DPEP of each target cell and pre-computed CMap DPEPs obtained from ^60^. Similarly, we obtained the scaled ranks of chemical compounds from DECCODE based on the reported similarity scores for comparison with other methods.

### Reproducibility analysis of CPS

To assess the reproducibility of CPS, we randomly subsampled 80% of cells in each cell type in both scRNA-seq and scATAC-seq atlases, and performed CPS calculations using these subsampled atlases. The sampling process was repeated ten times and the concordance of the CPS derived from the subsampled atlas data was quantified using Pearson’s correlation coefficient for all pairs of CPS. This procedure was applied to human and mouse data separately.

### Targeted TRN reconstruction after prediction

The reconstruction of the TRNs was performed using the igraph R package ^58^, incorporating multiple layers of molecular species and their interactions, from chemical compounds to kinases, kinases to TFs, and finally TFs to their target genes. The identification of chemical compound targets was confined to those fulfilling two criteria: 1) being targets of the given chemical compound in the curated database, and 2) being categorised as kinases. These kinases were then mapped to their potential targets which are themselves TFs. Interaction between the identified kinases and TFs was extracted from the STRING database. Lastly, the targets of the TFs were extracted from the TRRUST database. Nodes with positive CCS were included in the reconstructed TRNs.

### Gene set enrichment analysis

To evaluate if the CPS of genes computed by Refate prioritise genes that underlie cell identity, we performed over-representation based enrichment analysis using Fisher’s exact test on two gene sets including cell identity genes (CIGs) ^16^ and variably expressed TFs (VETFs) ^17^ using the top 1000 genes ranked by CPS. The top 1000 genes ranked by the SEG index were included as controls.

### Pathway enrichment analysis

To verify whether the CCS of genes calculated by Refate capture the molecular and cellular programs that underline cellular conversions, for each of the six cellular conversions, we performed pathway enrichment analysis on the Reactome and KEGG databases using the top 200 genes ranked by CCS and the *gost* function from the gprofiler2 package ^61^ with parameter ordered_query set to TRUE.

### Cell culture and neural crest differentiation

H9 (WiCell, WA09) human embryonic stem cells (hESC) were cultured in mTeSR Plus medium (StemCell Technologies, 100-0276) on matrigel-coated (Corning, 354277) six-well plates at 37°C in 20% O2 and 5% CO2. hESCs were passaged every 4–6 days at a split ratio of 1:50 to 1:100 using Versene (Life Technologies, 15040066) with media replenished daily. Cranial neural crest cells (CNCCs) were generated from H9-hESCs as previously described ^37,62^. H9 cells were cultured at 37°C in 5% O2 and 5% CO2. For single-cell dissociation, cells were treated with 0.5 mL of StemPro Accutase Cell Dissociation Reagent (Life Technologies, A1110501) per well of a 6-well plate and incubated at 37°C for 5 minutes. The dissociation was quenched with 2 mL of mTeSR Plus, and cells were triturated to ensure a uniform single-cell suspension. Cells were centrifuged at 1000 rpm for 5 minutes, resuspended in neural crest differentiation medium (DMEM/F12 (Life Technologies, 10565018) supplemented with 1 x serum-free B27 (Life Technologies, 17504044), 1 x Penicillin-Streptomycin (Life Technologies, 15140122), and counted using a hemocytometer. hESCs were seeded at a density of 3 × 10^4^ cells/cm² onto Matrigel-coated plates (µ-Plate 96 Well Round ibiTreat, #1.5 pol, Ibidi, 89606) and cultured in the neural crest differentiation medium. Y-27632 (StemCell Technologies, 72304) (10 µM) was added from day 0 to day 2 to promote cell survival and attachment. Differentiation was induced with CHIR99021 (3 µM) (Tocris, RDS442310) throughout the five-day protocol. Predicted small molecules (LDN193189, SB203580, Kenpaullone, Rimonabant, XL147, and Melatonin) were added to the media and were tested at three different concentrations (low, medium, and high, Figure 4A) to assess their impact on neural crest differentiation. Media with the compounds was replenished daily. The cells were cultured at 37°C in 5% O2 and 5% CO2.

### Immunohistochemistry

On day 5, cells were fixed with 4% paraformaldehyde (Scharlau, 30525-89-4) in DPBS (Thermo Fisher Scientific, 4190240) for 15 minutes, washed with PBS, and permeabilized with 0.1% Triton X-100 (Sigma, T8787) in PBS for 10 minutes. Blocking was performed with 1% bovine serum albumin (BSA) and 0.05% Tween-20 (Amresco, 0777) in PBS for 1 hour at room temperature. Cells were incubated overnight at 4°C with primary antibody Goat anti-SOX10 (1:200, R&D Systems, AF2864). The following day, cells were washed with PBS and incubated with Alexa Fluor-conjugated secondary antibodies (1:5000 dilution; Thermo Fisher Scientific) for 1 hour at room temperature. After washing with PBS, DAPI (1 µg/mL ThermoFisher Scientific, 62247) was used to stain nuclei for 5 minutes at room temperature. Finally cells were mounted with Ibidi Mounting Media (Ibidi, 50001-4).

Plates were imaged using an ImageXpress Confocal HT.ai High-Content Imaging System (Molecular Devices). For each well, at least five representative fields were captured at 20x magnification to ensure coverage of the entire well. Images were collected in two channels: DAPI for nuclear staining, FITC for SOX10 expression. Image preprocessing steps included background subtraction and correction for uneven illumination. Cell segmentation was performed on the DAPI channel to identify nuclei, followed by segmentation of the FITC channel to quantify SOX10 expression at the single-cell level. Cell segmentation was performed on the DAPI channel to identify nuclei, generating binary masks for single-cell regions. These masks were then applied to the FITC channel to quantify SOX10 expression at the single-cell level. Fluorescence intensity thresholds for identifying SOX10-positive cells were empirically determined and validated across multiple wells to ensure consistency and reproducibility.

SOX10 expression was analysed using several quantitative metrics. The number of SOX10-positive cells was determined by applying a fluorescence intensity threshold to the FITC channel. The proportion of SOX10-positive cells was calculated as the ratio of SOX10-positive cells to the total number of DAPI-stained nuclei. Mean SOX10 intensity per cell was computed by averaging fluorescence intensity across the segmented area of each cell. Total SOX10 intensity per image was determined by integrating the fluorescence signal across the entire field of view.

## Supplementary Figures

**Supplementary Figure 1.**
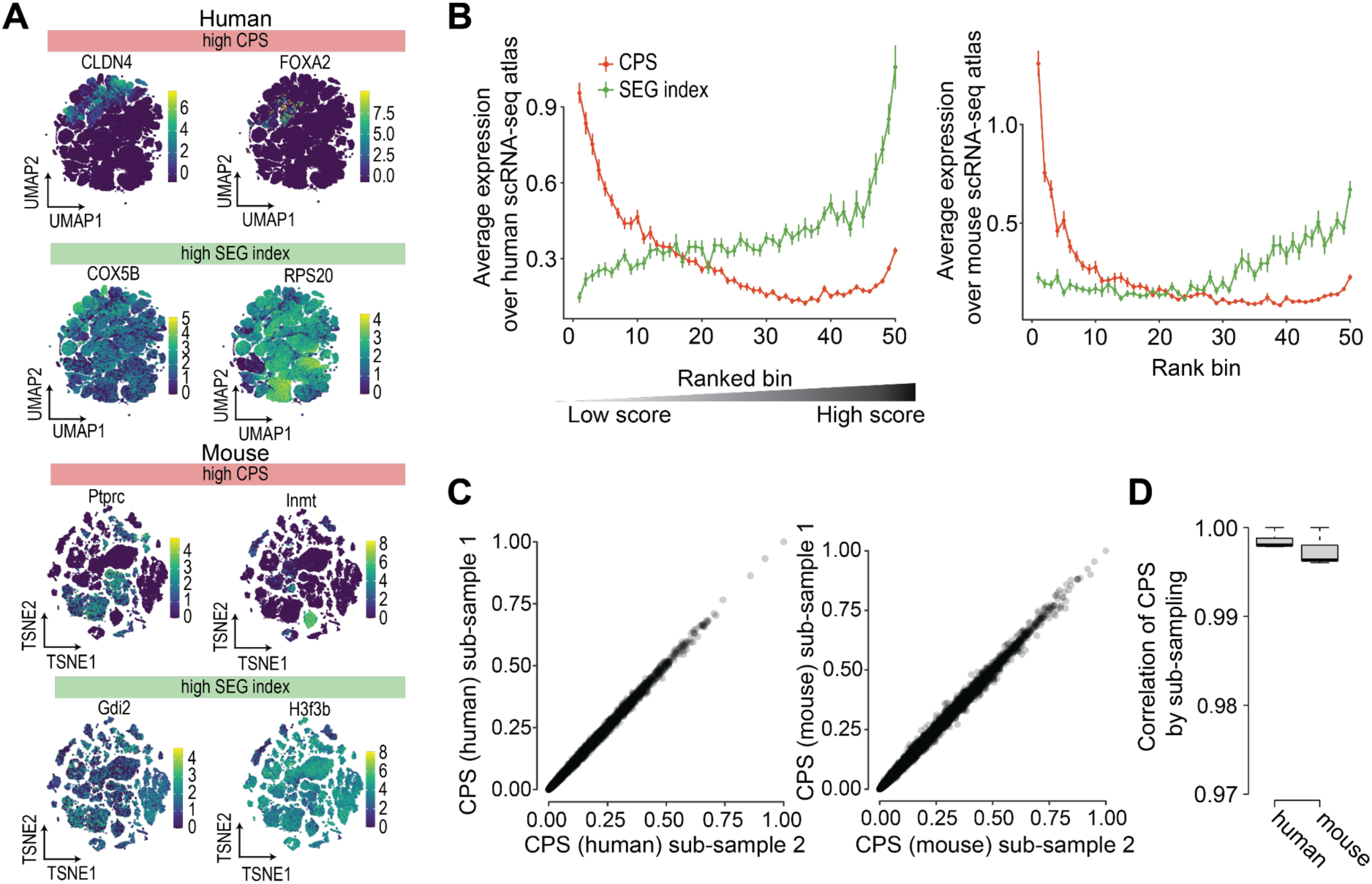
(related to Figure 1). Characterisation of cell propensity score (CPS). (**A**) Expression of representative genes with either high CPS or stably expressed gene (SEG) index in human and mouse scRNA-seq atlases. (**B**) Expression levels of genes ranked and binned either by CPS or SEG index for humans and mice, respectively. The expression level of each gene was calculated as the mean expression across all cells in the scRNA-seq atlas and the average of genes in each of the 50 bins are plotted on the y-axis. (**C, D**) Reproducibility assessment of CPS of genes derived from sub-sampling of 80% of cells from human and mouse atlases, respectively.

**Supplementary Figure 2.**
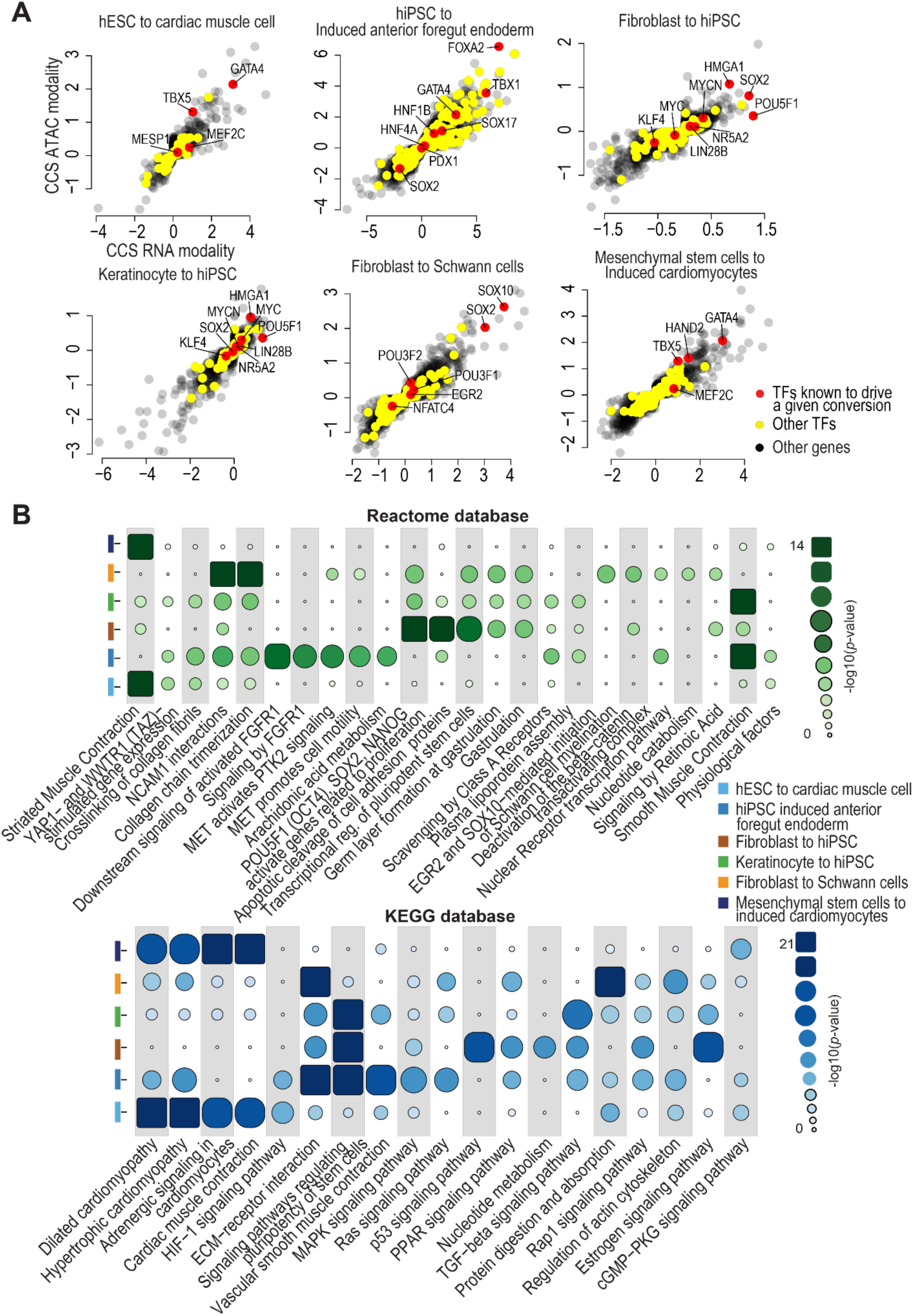
(related to Figure 2). Refate uncovers known and novel TFs that drive experimentally validated cellular conversions. (**A**) For each of the eight cellular conversions, scatterplots showing the CCS of known TFs (red), all other TFs (yellow), and all other genes (gray) computed on CPS derived from either RNA (x-axis) or ATAC (y-axis) modality, respectively. (**B**) Pathway enrichment analysis of the top-ranked genes by CCS for each cellular conversion using either Reactome or KEGG databases. The five most enriched pathways for each cellular conversion were shown.

**Supplementary Figure 3.**
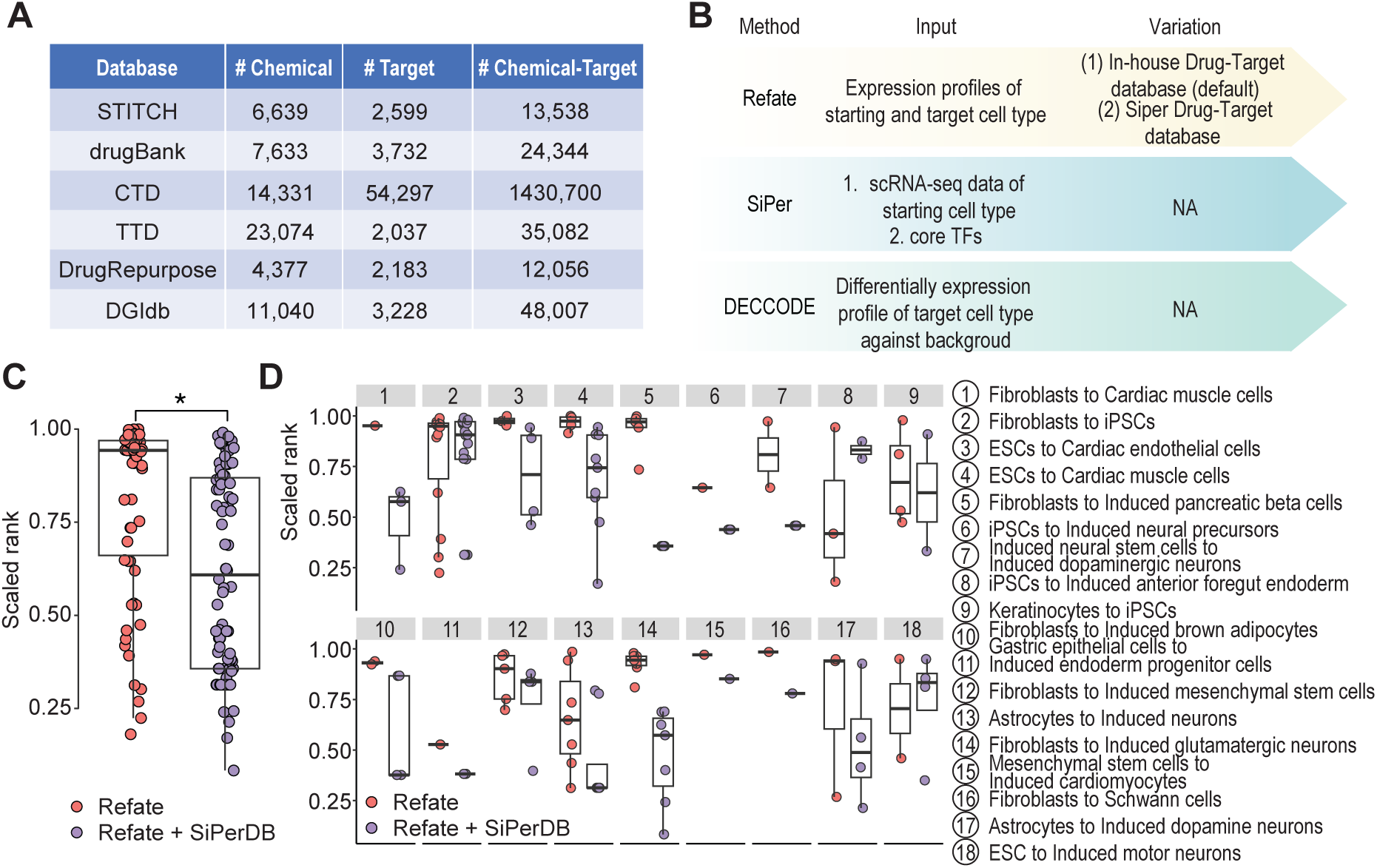
(related to Figure 3). Refate prioritises the known and novel chemical compounds for various cellular conversions. **(A)** Summary statistics of the six drug databases. **(B)** The benchmarking schematic of Refate, SiPer and DECCODE. For Refate, the drug-target database curated from this study or that curated from SiPer were tested. **(C)** Boxplots showing the overall scaled ranks of known chemical compounds for 18 curated cellular conversions by either Refate using drug-target database curated from this study or that from SiPer curated drug-target database. * denote statistical significance from a two-sided Wilcox rank sum test (p<0.05). **(D)** Breakdown of panel C to each of the 18 curated cellular conversions.

**Supplementary Figure 4.**
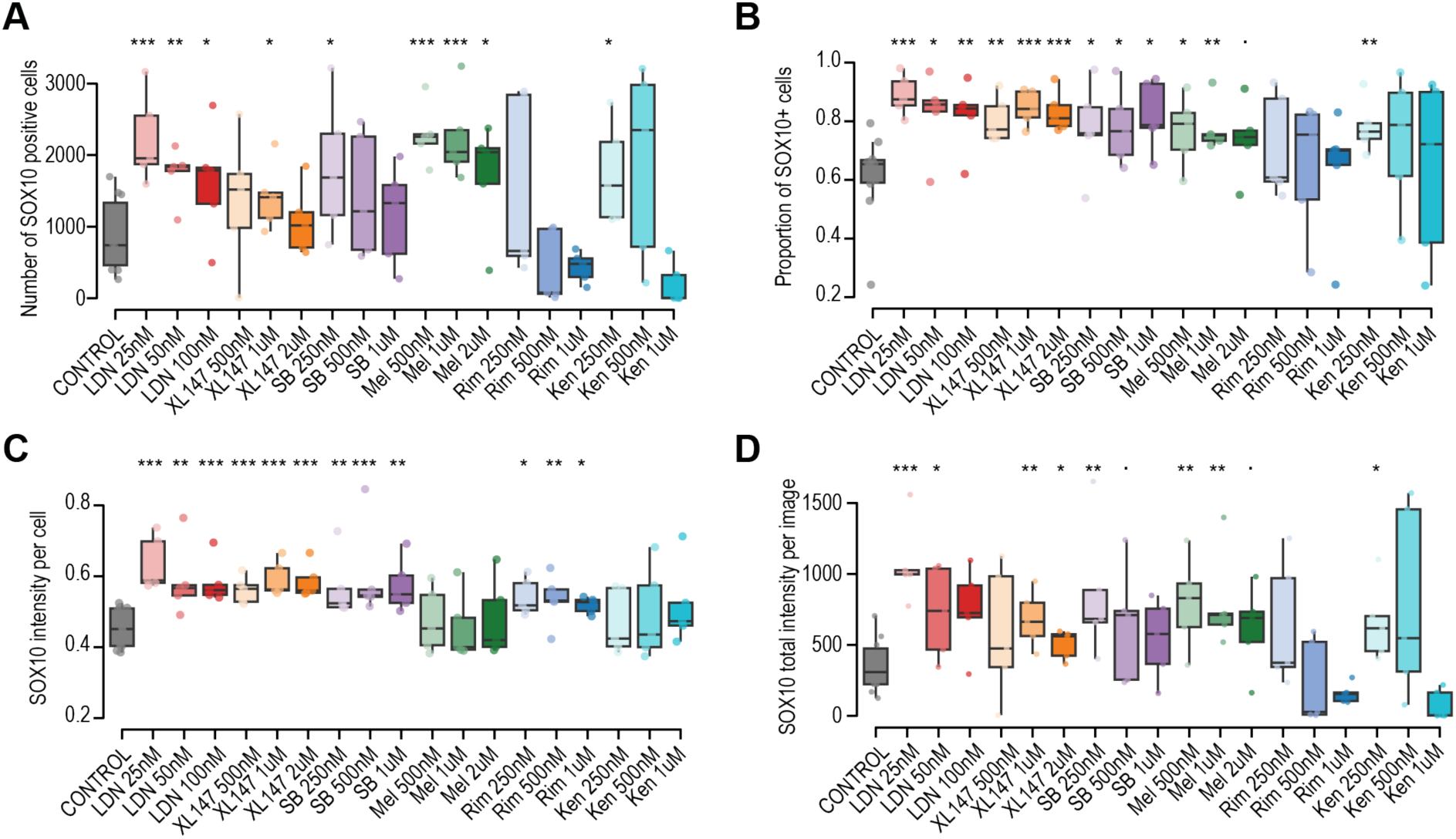
(related to Figure 4). Experimental validation of Refate-predicted small molecules for enhancing hESC to CNCC differentiation. Quantification of SOX10 expression across treatment conditions. Four key metrics were measured, including (**A**) the number of SOX10-positive cells, (**B**) the proportion of SOX10-positive cells (fraction of cells expressing SOX10), (**C**) SOX10 intensity per cell (mean fluorescence intensity per SOX10-positive cell), and (**D**) total SOX10 intensity per image (sum of fluorescence intensity in each image). One-way Wilcoxon rank sum test was performed. Asterisks above boxplots indicate statistically significant differences between those treatments against controls. Statistical significance: p < 0.1 (.), p < 0.05 (\**), p < 0.01 (*\*\*), p < 0.001 (***). Error bars indicate the standard deviation from five biological replicates.

## REFERENCES

1. Guo, C., and Morris, S.A. (2017). Engineering cell identity: establishing new gene regulatory and chromatin landscapes. Curr. Opin. Genet. Dev. 46, 50–57. 10.1016/j.gde.2017.06.011.

2. Morris, S.A., and Daley, G.Q. (2013). A blueprint for engineering cell fate: current technologies to reprogram cell identity. Cell Res. 23, 33–48. 10.1038/cr.2013.1.

3. Cahan, P., Cacchiarelli, D., Dunn, S.-J., Hemberg, M., Lopes, S.M.C. de S., Morris, S.A., Rackham, O.J.L., Sol, A. del, and Wells, C.A. (2021). Computational Stem Cell Biology: Open Questions and Guiding Principles. Cell Stem Cell 28, 20–32. 10.1016/j.stem.2020.12.012.

4. Tran, A., Yang, P., Yang, J.Y.H., and Ormerod, J. (2022). Computational approaches for direct cell reprogramming: from the bulk omics era to the single cell era. Brief. Funct. Genomics 21, 270–279. 10.1093/bfgp/elac008.

5. D’Alessio, A.C., Fan, Z.P., Wert, K.J., Baranov, P., Cohen, M.A., Saini, J.S., Cohick, E., Charniga, C., Dadon, D., Hannett, N.M., et al. (2015). A Systematic Approach to Identify Candidate Transcription Factors that Control Cell Identity. Stem Cell Rep. 5, 763–775. 10.1016/j.stemcr.2015.09.016.

6. Cahan, P., Li, H., Morris, S.A., da Rocha, E.L., Daley, G.Q., and Collins, J.J. (2014). CellNet: Network Biology Applied to Stem Cell Engineering. Cell 158, 903–915. 10.1016/j.cell.2014.07.020.

7. Rackham, O.J.L., Firas, J., Fang, H., Oates, M.E., Holmes, M.L., Knaupp, A.S., Suzuki, H., Nefzger, C.M., Daub, C.O., Shin, J.W., et al. (2016). A predictive computational framework for direct reprogramming between human cell types. Nat. Genet. 48, 331–335. 10.1038/ng.3487.

8. Xu, Q., Georgiou, G., Frölich, S., van der Sande, M., Veenstra, G.J.C., Zhou, H., and van Heeringen, S.J. (2021). ANANSE: an enhancer network-based computational approach for predicting key transcription factors in cell fate determination. Nucleic Acids Res. 49, 7966–7985. 10.1093/nar/gkab598.

9. Kamimoto, K., Stringa, B., Hoffmann, C.M., Jindal, K., Solnica-Krezel, L., and Morris, S.A. (2023). Dissecting cell identity via network inference and in silico gene perturbation. Nature, 1–10. 10.1038/s41586-022-05688-9.

10. Zheng, M., Xie, B., Okawa, S., Liew, S.Y., Deng, H., and Sol, A. del (2022). A single cell-based computational platform to identify chemical compounds targeting desired sets of transcription factors for cellular conversion. Stem Cell Rep. 10.1016/j.stemcr.2022.10.013.

11. Napolitano, F., Rapakoulia, T., Annunziata, P., Hasegawa, A., Cardon, M., Napolitano, S., Vaccaro, L., Iuliano, A., Wanderlingh, L.G., Kasukawa, T., et al. (2021). Automatic identification of small molecules that promote cell conversion and reprogramming. Stem Cell Rep. 16, 1381–1390. 10.1016/j.stemcr.2021.03.028.

12. Etchevers, H.C., Dupin, E., and Le Douarin, N.M. (2019). The diverse neural crest: from embryology to human pathology. Development 146, dev169821. 10.1242/dev.169821.

13. Kim, H.J., Wang, K., Chen, C., Lin, Y., Tam, P.P.L., Lin, D.M., Yang, J.Y.H., and Yang, P. (2021). Uncovering cell identity through differential stability with Cepo. Nat. Comput. Sci. 1, 784–790. 10.1038/s43588-021-00172-2.

14. Han, H., Cho, J.-W., Lee, S., Yun, A., Kim, H., Bae, D., Yang, S., Kim, C.Y., Lee, M., Kim, E., et al. (2018). TRRUST v2: an expanded reference database of human and mouse transcriptional regulatory interactions. Nucleic Acids Res. 46, D380–D386. 10.1093/nar/gkx1013.

15. Szklarczyk, D., Gable, A.L., Nastou, K.C., Lyon, D., Kirsch, R., Pyysalo, S., Doncheva, N.T., Legeay, M., Fang, T., Bork, P., et al. (2021). The STRING database in 2021: customizable protein–protein networks, and functional characterization of user-uploaded gene/measurement sets. Nucleic Acids Res. 49, D605–D612. 10.1093/nar/gkaa1074.

16. Xia, B., Zhao, D., Wang, G., Zhang, M., Lv, J., Tomoiaga, A.S., Li, Y., Wang, X., Meng, S., Cooke, J.P., et al. (2020). Machine learning uncovers cell identity regulator by histone code. Nat. Commun. 11, 2696. 10.1038/s41467-020-16539-4.

17. Shim, W.J., Sinniah, E., Xu, J., Vitrinel, B., Alexanian, M., Andreoletti, G., Shen, S., Sun, Y., Balderson, B., Boix, C., et al. (2020). Conserved Epigenetic Regulatory Logic Infers Genes Governing Cell Identity. Cell Syst. 11, 625–639.e13. 10.1016/j.cels.2020.11.001.

18. Y, L., S, G., D, S., A, W., E, P., Dm, L., T, S., Jyh, Y., and P, Y. (2019). Evaluating stably expressed genes in single cells. GigaScience 8. 10.1093/gigascience/giz106.

19. Consortium*, T.T.S., Jones, R.C., Karkanias, J., Krasnow, M.A., Pisco, A.O., Quake, S.R., Salzman, J., Yosef, N., Bulthaup, B., Brown, P., et al. (2022). The Tabula Sapiens: A multiple-organ, single-cell transcriptomic atlas of humans. Science. 10.1126/science.abl4896.

20. The Tabula Muris Consortium, Overall coordination, Logistical coordination, Organ collection and processing, Library preparation and sequencing, Computational data analysis, Cell type annotation, Writing group, Supplemental text writing group, and Principal investigators (2018). Single-cell transcriptomics of 20 mouse organs creates a Tabula Muris. Nature 562, 367–372. 10.1038/s41586-018-0590-4.

21. Zhang, K., Hocker, J.D., Miller, M., Hou, X., Chiou, J., Poirion, O.B., Qiu, Y., Li, Y.E., Gaulton, K.J., Wang, A., et al. (2021). A single-cell atlas of chromatin accessibility in the human genome. Cell 184, 5985–6001.e19. 10.1016/j.cell.2021.10.024.

22. Cusanovich, D.A., Hill, A.J., Aghamirzaie, D., Daza, R.M., Pliner, H.A., Berletch, J.B., Filippova, G.N., Huang, X., Christiansen, L., DeWitt, W.S., et al. (2018). A Single-Cell Atlas of In Vivo Mammalian Chromatin Accessibility. Cell 174, 1309–1324.e18. 10.1016/j.cell.2018.06.052.

23. Christoforou, N., Chakraborty, S., Kirkton, R.D., Adler, A.F., Addis, R.C., and Leong, K.W. (2017). Core Transcription Factors, MicroRNAs, and Small Molecules Drive Transdifferentiation of Human Fibroblasts Towards The Cardiac Cell Lineage. Sci. Rep. 7, 40285. 10.1038/srep40285.

24. Takahashi, K., and Yamanaka, S. (2006). Induction of Pluripotent Stem Cells from Mouse Embryonic and Adult Fibroblast Cultures by Defined Factors. Cell 126, 663–676. 10.1016/j.cell.2006.07.024.

25. Santra, T., and Delatola, E.I. (2016). A Bayesian algorithm for detecting differentially expressed proteins and its application in breast cancer research. Sci. Rep. 6, 30159. 10.1038/srep30159.

26. Heng, J.-C.D., Feng, B., Han, J., Jiang, J., Kraus, P., Ng, J.-H., Orlov, Y.L., Huss, M., Yang, L., Lufkin, T., et al. (2010). The Nuclear Receptor Nr5a2 Can Replace Oct4 in the Reprogramming of Murine Somatic Cells to Pluripotent Cells. Cell Stem Cell 6, 167–174. 10.1016/j.stem.2009.12.009.

27. Kuo, I.Y., and Ehrlich, B.E. (2015). Signaling in Muscle Contraction. Cold Spring Harb. Perspect. Biol. 7, a006023. 10.1101/cshperspect.a006023.

28. Mattingly, C.J., Rosenstein, M.C., Davis, A.P., Colby, G.T., Forrest, J.N., and Boyer, J.L. (2006). The Comparative Toxicogenomics Database (CTD): A Cross-Species Resource for Building Chemical-Gene Interaction Networks. Toxicol. Sci. Off. J. Soc. Toxicol. 92, 587–595. 10.1093/toxsci/kfl008.

29. Freshour, S.L., Kiwala, S., Cotto, K.C., Coffman, A.C., McMichael, J.F., Song, J.J., Griffith, M., Griffith, O.L., and Wagner, A.H. (2021). Integration of the Drug–Gene Interaction Database (DGIdb 4.0) with open crowdsource efforts. Nucleic Acids Res. 49, D1144–D1151. 10.1093/nar/gkaa1084.

30. Wishart, D.S., Feunang, Y.D., Guo, A.C., Lo, E.J., Marcu, A., Grant, J.R., Sajed, T., Johnson, D., Li, C., Sayeeda, Z., et al. (2018). DrugBank 5.0: a major update to the DrugBank database for 2018. Nucleic Acids Res. 46, D1074–D1082. 10.1093/nar/gkx1037.

31. Corsello, S.M., Bittker, J.A., Liu, Z., Gould, J., McCarren, P., Hirschman, J.E., Johnston, S.E., Vrcic, A., Wong, B., Khan, M., et al. (2017). The Drug Repurposing Hub: a next-generation drug library and information resource. Nat. Med. 23, 405–408. 10.1038/nm.4306.

32. Kuhn, M., von Mering, C., Campillos, M., Jensen, L.J., and Bork, P. (2008). STITCH: interaction networks of chemicals and proteins. Nucleic Acids Res. 36, D684–D688. 10.1093/nar/gkm795.

33. Zhu, F., Han, B., Kumar, P., Liu, X., Ma, X., Wei, X., Huang, L., Guo, Y., Han, L., Zheng, C., et al. (2010). Update of TTD: Therapeutic Target Database. Nucleic Acids Res. 38, D787–D791. 10.1093/nar/gkp1014.

34. Zhu, S., Li, W., Zhou, H., Wei, W., Ambasudhan, R., Lin, T., Kim, J., Zhang, K., and Ding, S. (2010). Reprogramming of Human Primary Somatic Cells by OCT4 and Chemical Compounds. Cell Stem Cell 7, 651–655. 10.1016/j.stem.2010.11.015.

35. Simões-Costa, M., and Bronner, M.E. (2015). Establishing neural crest identity: a gene regulatory recipe. Dev. Camb. Engl. 142, 242–257. 10.1242/dev.105445.

36. Zyner, K.G., Simeone, A., Flynn, S.M., Doyle, C., Marsico, G., Adhikari, S., Portella, G., Tannahill, D., and Balasubramanian, S. (2022). G-quadruplex DNA structures in human stem cells and differentiation. Nat. Commun. 13, 142. 10.1038/s41467-021-27719-1.

37. Gomez, G.A., Prasad, M.S., Sandhu, N., Shelar, P.B., Leung, A.W., and García-Castro, M.I. (2019). Human Neural Crest Induction by Temporal Modulation of WNT Activation. Dev. Biol. 449, 99–106. 10.1016/j.ydbio.2019.02.015.

38. Asih, P.R., Prikas, E., Stefanoska, K., Tan, A.R.P., Ahel, H.I., and Ittner, A. (2020). Functions of p38 MAP Kinases in the Central Nervous System. Front. Mol. Neurosci. 13. 10.3389/fnmol.2020.570586.

39. Wei, Z.Z., Yu, S.P., Lee, J.H., Chen, D., Taylor, T.M., Deveau, T.C., Yu, A.C.H., and Wei, L. (2014). Regulatory Role of the JNK-STAT1/3 Signaling in Neuronal Differentiation of Cultured Mouse Embryonic Stem Cells. Cell. Mol. Neurobiol. 34, 881– 893. 10.1007/s10571-014-0067-4.

40. Foshay, K.M., and Gallicano, G.I. (2008). Regulation of Sox2 by STAT3 Initiates Commitment to the Neural Precursor Cell Fate. Stem Cells Dev. 17, 269–278. 10.1089/scd.2007.0098.

41. FitzPatrick, L.M., Hawkins, K.E., Delhove, J.M.K.M., Fernandez, E., Soldati, C., Bullen, L.F., Nohturfft, A., Waddington, S.N., Medina, D.L., Bolaños, J.P., et al. (2018). NF-κB Activity Initiates Human ESC-Derived Neural Progenitor Cell Differentiation by Inducing a Metabolic Maturation Program. Stem Cell Rep. 10, 1766–1781. 10.1016/j.stemcr.2018.03.015.

42. Feng, Z., and Porter, A.G. (1999). NF-κB/Rel Proteins Are Required for Neuronal Differentiation of SH-SY5Y Neuroblastoma Cells *. J. Biol. Chem. 274, 30341–30344. 10.1074/jbc.274.43.30341.

43. Accili, D., and Arden, K.C. (2004). FoxOs at the Crossroads of Cellular Metabolism, Differentiation, and Transformation. Cell 117, 421–426. 10.1016/S0092-8674(04)00452-0.

44. Nerlov, C. (2007). The C/EBP family of transcription factors: a paradigm for interaction between gene expression and proliferation control. Trends Cell Biol. 17, 318–324. 10.1016/j.tcb.2007.07.004.

45. EGF–ERBB signalling: towards the systems level | Nature Reviews Molecular Cell Biology https://www.nature.com/articles/nrm1962.

46. Saxton, R.A., and Sabatini, D.M. (2017). mTOR Signaling in Growth, Metabolism, and Disease. Cell 168, 960–976. 10.1016/j.cell.2017.02.004.

47. Polo, J.M., Anderssen, E., Walsh, R.M., Schwarz, B.A., Nefzger, C.M., Lim, S.M., Borkent, M., Apostolou, E., Alaei, S., Cloutier, J., et al. (2012). A Molecular Roadmap of Reprogramming Somatic Cells into iPS Cells. Cell 151, 1617–1632. 10.1016/j.cell.2012.11.039.

48. Ng, A.H.M., Khoshakhlagh, P., Rojo Arias, J.E., Pasquini, G., Wang, K., Swiersy, A., Shipman, S.L., Appleton, E., Kiaee, K., Kohman, R.E., et al. (2021). A comprehensive library of human transcription factors for cell fate engineering. Nat. Biotechnol. 39, 510–519. 10.1038/s41587-020-0742-6.

49. Joung, J., Ma, S., Tay, T., Geiger-Schuller, K.R., Kirchgatterer, P.C., Verdine, V.K., Guo, B., Arias-Garcia, M.A., Allen, W.E., Singh, A., et al. (2023). A transcription factor atlas of directed differentiation. Cell 186, 209–229.e26. 10.1016/j.cell.2022.11.026.

50. Parekh, U., Wu, Y., Zhao, D., Worlikar, A., Shah, N., Zhang, K., and Mali, P. (2018). Mapping Cellular Reprogramming via Pooled Overexpression Screens with Paired Fitness and Single-Cell RNA-Sequencing Readout. Cell Syst. 7, 548–555.e8. 10.1016/j.cels.2018.10.008.

51. Kim, H.J., Kim, T., Hoffman, N.J., Xiao, D., James, D.E., Humphrey, S.J., and Yang, P. (2021). PhosR enables processing and functional analysis of phosphoproteomic data. Cell Rep. 34, 108771.

52. Allaire, J., Ellis, P., Gandrud, C., Kuo, K., Lewis, B., Owen, J., Russell, K., Rogers, J., Sese, C., Yetman, C., et al. (2017). Package ‘networkD3.’ D3 JavaScript Netw. Graphs R.

53. Schindelin, J., Arganda-Carreras, I., Frise, E., Kaynig, V., Longair, M., Pietzsch, T., Preibisch, S., Rueden, C., Saalfeld, S., Schmid, B., et al. (2012). Fiji: an open-source platform for biological-image analysis. Nat. Methods 9, 676–682. 10.1038/nmeth.2019.

54. Stirling, D.R., Swain-Bowden, M.J., Lucas, A.M., Carpenter, A.E., Cimini, B.A., and Goodman, A. (2021). CellProfiler 4: improvements in speed, utility and usability. BMC Bioinformatics 22, 433. 10.1186/s12859-021-04344-9.

55. Pliner, H.A., Packer, J.S., McFaline-Figueroa, J.L., Cusanovich, D.A., Daza, R.M., Aghamirzaie, D., Srivatsan, S., Qiu, X., Jackson, D., Minkina, A., et al. (2018). Cicero Predicts cis-Regulatory DNA Interactions from Single-Cell Chromatin Accessibility Data. Mol. Cell 71, 858–871.e8. 10.1016/j.molcel.2018.06.044.

56. Nguyen, Q.H., Lukowski, S.W., Chiu, H.S., Senabouth, A., Bruxner, T.J.C., Christ, A.N., Palpant, N.J., and Powell, J.E. (2018). Single-cell RNA-seq of human induced pluripotent stem cells reveals cellular heterogeneity and cell state transitions between subpopulations. Genome Res. 28, 1053–1066. 10.1101/gr.223925.117.

57. Churko, J.M., Garg, P., Treutlein, B., Venkatasubramanian, M., Wu, H., Lee, J., Wessells, Q.N., Chen, S.-Y., Chen, W.-Y., Chetal, K., et al. (2018). Defining human cardiac transcription factor hierarchies using integrated single-cell heterogeneity analysis. Nat. Commun. 9, 4906. 10.1038/s41467-018-07333-4.

58. Csardi, G., and Nepusz, T. (2006). The igraph software package for complex network research. InterJournal Complex Syst. 1695, 1–9.

59. Subramanian, A., Narayan, R., Corsello, S.M., Peck, D.D., Natoli, T.E., Lu, X., Gould, J., Davis, J.F., Tubelli, A.A., Asiedu, J.K., et al. (2017). A Next Generation Connectivity Map: L1000 platform and the first 1,000,000 profiles. Cell 171, 1437–1452.e17. 10.1016/j.cell.2017.10.049.

60. Napolitano, F., Carrella, D., Gao, X., and di Bernardo, D. (2020). gep2pep: a bioconductor package for the creation and analysis of pathway-based expression profiles. Bioinformatics 36, 1944–1945. 10.1093/bioinformatics/btz803.

61. Kolberg, L., Raudvere, U., Kuzmin, I., Vilo, J., and Peterson, H. (2020). gprofiler2--an R package for gene list functional enrichment analysis and namespace conversion toolset g:Profiler. F1000Research 9, ELIXIR-709. 10.12688/f1000research.24956.2.

62. Wilderman, A., D’haene, E., Baetens, M., Yankee, T.N., Winchester, E.W., Glidden, N., Roets, E., Van Dorpe, J., Janssens, S., Miller, D.E., et al. (2024). A distant global control region is essential for normal expression of anterior HOXA genes during mouse and human craniofacial development. Nat. Commun. 15, 136. 10.1038/s41467-023-44506-2.

